# T cell cholesterol transport is a metabolic checkpoint that links intestinal immune responses to dietary lipid absorption

**DOI:** 10.1101/2024.03.08.584164

**Authors:** Yajing Gao, John P. Kennelly, Xu Xiao, Emily Whang, Alessandra Ferrari, Alexander H. Bedard, Julia J. Mack, Alexander H. Nguyen, Thomas Weston, Lauren F. Uchiyama, Min Sub Lee, Stephen G. Young, Steven J. Bensinger, Peter Tontonoz

## Abstract

The intrinsic pathways that control membrane organization in immune cells and the impact of such pathways on cellular function are not well defined. Here we report that the non-vesicular cholesterol transporter Aster-A links plasma membrane (PM) cholesterol availability in T cells to immune signaling and systemic metabolism. Aster-A is recruited to the PM during T-cell receptor (TCR) activation, where it facilitates the removal of newly generated “accessible” membrane cholesterol. Loss of Aster-A leads to excess PM cholesterol accumulation, resulting in enhanced TCR nano-clustering and signaling, and Th17 cytokine production. Finally, we show that the mucosal Th17 response is restrained by PM cholesterol remodeling. Ablation of Aster-A in T cells leads to enhanced IL-22 production, reduced intestinal fatty acid absorption, and resistance to diet-induced obesity. These findings delineate a multi-tiered regulatory scheme linking immune cell lipid flux to nutrient absorption and systemic physiology.

## Introduction

The PM is the major cellular reserve for cholesterol. The PM is thought to contain three cholesterol pools of distinct biochemical and biophysical properties (*1*): 1. an essential pool required for membrane integrity and cell viability; 2. a relatively inactive pool (recognized by Ostreolysin A/OlyA) in which cholesterol is sequestered by sphingomyelin (SM) in nanodomains (*2*), and 3. an ‘active/accessible’ pool that can be readily mobilized (marked by accessibility to the domain 4 of Anthrolysin O/ALOD4 probe (*3*, *4*). Given its important role in cellular homeostasis several recent studies have focused on cellular and external cues that regulate the size of the accessible cholesterol, such as uptake of lipoprotein-derived cholesterol, activation of sphingomyelin degradation, and exposure to infection-elicited defense molecules (*5*– *11*). Potential fates of excess PM accessible cholesterol include movement to the ER for esterification or efflux to extracellular lipoprotein acceptors (*6*, *12–15*). However, the roles of this bioactive pool in PM itself are poorly defined. One exception is hedgehog signaling at the primary cilium, where PTCH1 inactivation by hedgehog ligands increases accessible cholesterol which subsequently potentiate the activation of Smoothened (*8*, *16*).

Non-vesicular sterol transport is now recognized to be important for cellular lipid homeostasis and maintaining appropriate gradients between membrane compartments (*17–19*). The Aster proteins play an indispensable role in the movement of accessible cholesterol from PM to ER (*5*, *12*, *20*). Increased accessible PM cholesterol facilitates the recruitment of the GRAM domain of Asters to PM and the formation of PM-ER contact sites, through which the ASTER domain channels excess cholesterol to the ER (*12*). Aster family proteins are required for the efficient internalization, storage, and utilization of diet- and lipoprotein-derived cholesterol in various metabolic and steroidogenic organs (*6*, *12*, *15*, *21*). Their function in other cell types remain to be elucidated. Of particular interest, Aster-A is highly expressed various lymphoid organs, suggesting an unrecognized function for sterol transport in the immune system.

Studies using chemical or indirect genetic approaches levels have reported that increasing PM abundance can enhance T cell receptor (TCR) signaling (*22–26*). At the same time, recent cryo-EM structural studies have revealed that the resting αβTCR core contains two free cholesterol molecules that may serve to maintain the inactivate state (*27–30*). Such studies imply that PM cholesterol availability must be tightly controlled during immune activation. However, it remains unknown whether T cells engage specific machinery to monitor and redistribute PM cholesterol. How membrane cholesterol dynamics are integrated with TCR signaling cascades has also not been defined.

Here we show that non-vesicular cholesterol transport restrains intestinal Th17 responses by controlling PM cholesterol availability and TCR nanoclustering. We further reveal that loss of Aster-A in T cells causes dietary fatty acid malabsorption and resistance to diet induced obesity. Microbiota-dependent IL-22 production from Aster-A-deficient intestinal Th17 cells suppresses fatty acid uptake by the small intestine. These findings define an immunoregulatory role of intracellular lymphocyte cholesterol trafficking and uncover the importance of Aster-A in maintaining mucosal homeostasis and intestinal lipid metabolism.

## Results

### Nonvesicular cholesterol transport in T lymphocytes is required for optimal dietary fatty acid absorption

We discovered that whole-body deletion of the nonvesicular cholesterol transporter Aster-A profoundly impaired fatty acid uptake in the small intestine (SI) (**Fig. 1a**). Aster-A-deficient (A^-/-^) mice exhibited reduced maximum [^3^H] uptake in jejunum and reduced total SI [^3^H] level following gavage with [^3^H]-triolein (**Fig. S1a**). There was also a delay in the appearance of [^3^H] in plasma of mice lacking Aster-A compared to controls (**Fig. S1b**).

**Figure 1.**
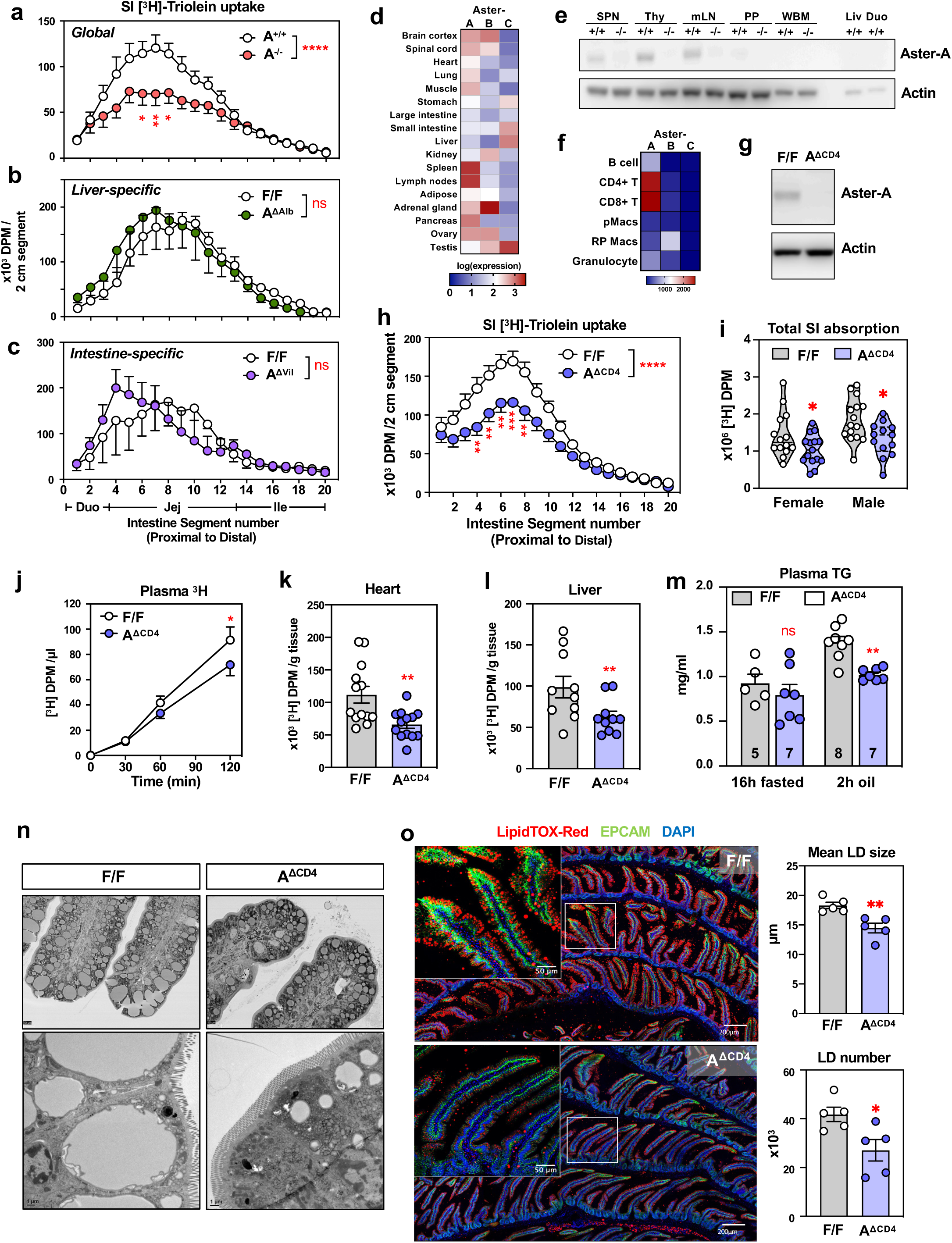
Aster-A deficiency impairs fatty acid absorption in the small intestine. **a.** Distribution of radioactivity in intestinal segments of Aster-A wiltype (A^+/+^, n=13) and global deficient (A^-/-^, n=14) littermate mice after an oral challenge of olive oil containing [^3^H]-triolein (18:0) for 2 h. **b-c.** Distribution of radioactivity in intestinal segments of liver/hepatocyte-specific (A^ΔAlb^,**b**), or intestinal epithelium-specific (A^ΔVil^, **c**) knockout mice and their cre-negative floxed (F/F) littermate control after an oral challenge of olive oil containing [^3^H]-triolein (18:0) for 2 h (n=9 and 6 /group for liver-specific knockout mice; n=4 /group for intestinal epithelium-specific mice). **d.** Expression of Aster-A, B, C (transcript *Gramd1a, Gramd1b, Gramd1c*) in mouse tissues from BioGPS dataset. **e.** Western blot analysis of expression of Aster-A in the indicated tissues from Aster-A global deficient (A^-/-^) or wiltype (A^+/+^) littermate. SPN, spleen; Thy, Thymus; mLN, mesenteric lymph nodes; PP, Peyer’s patches; WBM, whole bone marrow; Liv, liver; Duo, duodenum. Equal quantity of total protein (10μg) was used in each lane. **f.** Expression of Aster-A, B, C in mouse immune populations from the Immgen database. **g.** Western blot analysis of expression of Aster-A in T cells from Aster-A^fl/fl^ (F/F) and littermates Aster-A^fl/fl^ CD4-Cre (A^ΔCD4^) mice. **h.** Distribution of radioactivity in intestinal segments of male and female Aster-A^fl/fl^ (F/F, n=29) and littermates Aster-A^fl/fl^ CD4-Cre (A^ΔCD4^, n=28) mice after an oral challenge of olive oil containing [^3^H]-triolein (18:0) for 2 h. **i.** Total radioactivity in small intestines from experiments in **h**, separated by sex (Female, n=14/15 per group; Male, n=15/13 per group). **j.** Kinetics of radioactivity in plasma of Aster-A^fl/fl^ (F/F, n=25) and littermates Aster-A^fl/fl^ CD4-Cre (A^ΔCD4^, n=23) after an oral challenge of olive oil containing [^3^H]-triolein (18:0). **k-l.** Radioactivity in heart (**k,** n=13 /group) and liver (**l,** n=10 /group) from Aster-A^fl/fl^ (F/F) and littermates Aster-A^fl/fl^ CD4-Cre (A^ΔCD4^) after an oral challenge of olive oil containing [^3^H]-triolein (18:0) for 2 h. **m.** Non-radioactive plasma triglyceride level from female Aster-A^fl/fl^ (F/F) and littermates Aster-A^fl/fl^ CD4-Cre (A^ΔCD4^) after 16 h fasting or 16 h fasting followed by an oral challenge of olive oil for 2 h. **n.** Electron microscopy images of proximal jejunum villi tip (above) and brush border (below) from Aster-A^fl/fl^ (F/F) and littermates Aster-A^fl/fl^ CD4-Cre (A^ΔCD4^) after an oral challenge of olive oil for 2 h. **o.** Visualization of neutral lipid accumulation (marked by LipidTOX-Red) in jejunum cross sections from Aster-A^fl/fl^ (F/F) and littermates Aster-A^fl/fl^ CD4-Cre (A^ΔCD4^) mice after an oral challenge of olive oil for 2 h. Scale bar: 200μm. Inlay shows villi tip in higher magnification. Representative of 5 mice/group. Scale bar: 50μm. Right, mean lipid droplet (LD) size (in μm) and total number of LD per image. n=5 mice /group. Statistical analysis: for panels a, b, c, h, m, two-way ANOVA with Sidak’s multiple comparison test. For panel i, mixed-effects analysis. For panel j, Repeated Measure two-way ANOVA with Sidak’s multiple comparison test. For panel k, l, o, two-tailed unpaired Welch’s t-test. ns, p>0.05, *p<0.05, **p<0.01, ***p<0.001, ****p<0.0001.

We next sought to pinpoint the tissue specific basis of this effect. We have previously shown that Aster proteins play important roles in enterohepatic handling of dietary cholesterol. In liver, Aster-A and Aster-C together facilitate HDL-cholesterol uptake and biliary excretion (*6*). In the intestinal epithelium, Aster-B and Aster-C are required for dietary cholesterol uptake (*15*). However, Aster-A mRNA expression in SI is 26.3- and 7.6-fold lower than Aster-B and Aster-C respectively (*15*), and it does not respond to dietary fat intake (**Fig. S1c**). To determine whether loss of Aster-A alone in the enterohepatic compartment was sufficient to impact dietary fatty acid uptake, we deleted Aster-A in liver (by Albumin-Cre, referred to as A^ΔAlb^ mice) or intestinal epithelium (by Villin-CreERT, referred to as A^ΔVil^ mice) of mice (**Fig. S1d**). Neither A^ΔAlb^ nor A^ΔVil^ mice showed a difference in fatty acid absorption compared to their respective littermate controls (**Fig. 1b-c**). Thus, the absorption defect of Aster-A global KO mice is not due to intrinsic effects in enterohepatic tissues.

We then sought to identify tissues and/or cell types that have notable expression of Aster-A, but not Aster-B or Aster-C. In public mRNA expression datasets from humans and mice, secondary lymphoid organs (spleen and lymph nodes) exhibit abundant transcripts encoding Aster-A (*GRAMD1A*/*Gramd1a*), and low to trace levels of those encoding Aster-B and -C (**Fig. 1d-e**). Examination of specific immune cell types in the Immgen database further revealed high expression of Aster-A in CD4 and CD8 T cells compared to other immune cell types such as macrophages, which we have shown previously to express predominantly Aster-B (**Fig. 1f**) (*12*). To test if T cell- specific Aster-A expression influenced SI fat absorption, we generated mice lacking Aster-A in conventional T cells using CD4-Cre (referred to as Α^ΔCD4^ mice). A^ΔCD4^ mice showed more than a 100-fold reduction in *Gramd1a* transcripts in CD4+ and CD8+ T cells and undetectable Aster-A protein (**Fig. 1g and S1e**). A^ΔCD4^ mice were born at Mendelian ratios and showed normal growth curves from weaning to early adulthood (**Fig. S1f,g**), suggesting that T-cell-specific deletion of Aster-A did not grossly impact development or lead to early onset immune disorders.

To examine whether T cell-intrinsic Aster-A impacted dietary fatty acid uptake, we gavaged A^ΔCD4^ or littermate control floxed mice with [^3^H]-labeled triolein and evaluated appearance of radioactivity in SI segments, plasma, and tissues. A^ΔCD4^ mice showed reduced peak [^3^H] uptake in distal duodenum to jejunum and reduced total intestinal [^3^H] absorption (**Fig. 1h-i**). Interestingly, the magnitude of the defect was comparable to that observed in Aster-A global KO animals (**Fig. 1a**). A^ΔCD4^ mice showed a reduced rate of appearance of label in plasma, as well as lower [^3^H] tissue uptake in heart and Liver (**Fig.1j-l**). Despite an overall higher fatty acid absorptive capacity in male mice, impaired fatty acid was observed in both A^ΔCD4^ mice of both sexes (**Fig. 1i and S1h**). In line with our tracer studies, an oral challenge of unlabeled olive oil also gave rise to lower total plasma triglyceride levels in A^ΔCD4^ mice after 2 h (**Fig. 1m**). Thus, T-cell specific Aster-A deletion reduces fatty acid uptake into intestinal epithelium, slows the subsequent entry of dietary fatty acids into circulation, and impairs their tissue uptake.

Of note, we found no difference in the length of small intestine between control and A^ΔCD4^ mice fed chow diet (**Fig.S1i**). Similar total gastrointestinal transit time was also observed between genotypes, suggesting peristalsis was not impaired (**Fig.S1j**). Histologically, jejunal villi and crypt morphology, villi length, and brush border morphology of A^ΔCD4^ mice were indistinguishable from control littermates (**Fig.1n and S1k**). We found no histological signs of overt inflammation in the lamina propria (**Fig. S1k**). Following olive oil gavage, enterocytes at villi tip of control mice accumulated large, unilocular lipid droplets, whereas enterocytes of A^ΔCD4^ contained markedly smaller lipid droplets (**Fig. 1n and S1k**). To quantify the distribution of enterocyte lipid droplets in an unbiased manner, we fluorescently labeled neutral lipids, imaged a large cross section of jejunum, and analyzed lipid content by CellProfiler (**Fig.1o**). A^ΔCD4^ mice exhibited a major shift in distribution to smaller droplet size and reduced total numbers of lipid droplets (**Fig. 1o and S1m**). This finding was accompanied by reduced expression of lipid droplet-associated gene expression in jejunal epithelium as assessed by RNA-seq (**Fig. S1n**). No impairment was observed in oral glucose absorption (**Fig. S1l**). Thus, the impact of Aster-A in T cells on nutrient absorption is specific to fatty acid uptake and unrelated to any apparent SI pathology.

To confirm that the defect in fatty acid uptake in A^ΔCD4^ mice was not due to the expression of CD4^Cre^, we generated mice that harbored the CD4^Cre^ allele but were heterozygous for the floxed Aster-A allele (CD4^Cre^; A^fl/+^). We found no difference in [^3^H]- triolein absorption between these mice and Cre-negative; A^fl/+^ littermates (**Fig.S1o-r**). This finding confirmed that defective fatty acid absorption in A^ΔCD4^ mice was linked to loss of Aster-A function in T cells.

### Loss of T-cell-specific Aster-A prevents diet-induced obesity

To understand the long-term metabolic impact of Aster-A loss in T cells, we challenged mice with a 60 kcal% high-fat diet (HFD). Compared to littermate controls, both female and male A^ΔCD4^ mice were resistant to HFD-induced weight gain (**Fig. 2a**). After 10 weeks of HFD feeding, they were indistinguishable from controls in body length, but phenotypically leaner (**Fig. 2b**). Reductions in tissue weights of liver, eWAT, and iWAT were also observed in A^ΔCD4^ mice (**Fig. 2c and S2a**). Analysis of body composition by MRI showed reduced total fat mass in A^ΔCD4^ mice but no difference in lean mass (**Fig. 2d and S2b**). There were no body composition differences between genotypes when mice were fed normal chow (4 kcal% fat) (**Fig. S2c**), implying that T-cell specific loss of Aster-A impairs the maximum fatty acid handling capacity of the SI epithelium.

**Figure 2.**
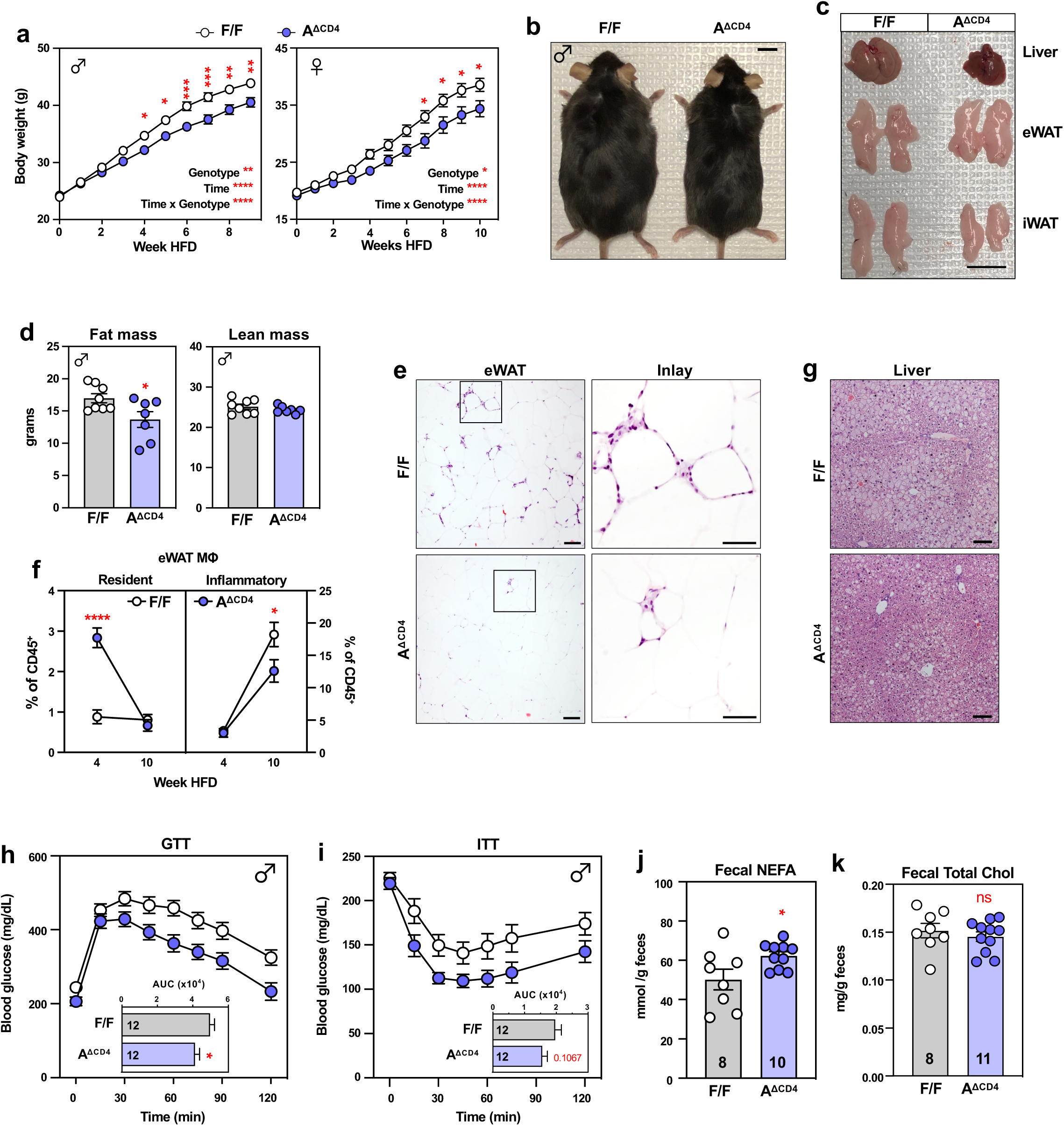
Lymphocyte non-vesicular cholesterol transport regulates diet-induced obesity. **a.** Body weight of male (left) and female (right) Aster-A^fl/fl^ (F/F) and littermates Aster-A^fl/fl^ CD4-Cre (A^ΔCD4^) mice fed ad libitum with 60 kcal% high fat diet (HFD) from 8-10 weeks of age for 9-10 weeks. Male, n=17 (F/F) and 16 (A^ΔCD4^); Female, n=15/group. **b-c.** Representative of external (**b**) and gross tissue (**c**) appearance of male Aster-A^fl/fl^ (F/F) and littermates Aster-A^fl/fl^ CD4-Cre (A^ΔCD4^) mice after 10-weeks of HFD feeding. **d.** Fat and lean mass of male mice in (**a-c**) determined by EchoMRI. n=8 (F/F) and 7 (A^ΔCD4^). **e.** Representative hematoxylin and eosin (H&E) histology of epididymal WAT from mice in (**a**). Scale bar, 100μm; Inlay scale bar, 50μm. **f.** Percentage of tissue-resident (F4/80^hi^ CD11b^+^) and inflammatory (F4/80^+^CD11b^hi^) macrophages in eWAT of Aster-A^fl/fl^ (F/F) and littermates Aster-A^fl/fl^ CD4-Cre (A^ΔCD4^) mice fed *ad libitum* with HFD for 4 or 10 weeks. n=3-5/group. **g.** Representative hematoxylin and eosin (H&E) histology of Liver from mice in (**a**). Scale bar, 100μm; **h-i.** intraperitoneal glucose tolerance test (GTT, **h**) and insulin tolerance test (ITT, **i**) on Aster-A^fl/fl^ (F/F) and littermates Aster-A^fl/fl^ CD4-Cre (A^ΔCD4^) mice fed *ad libitum* with HFD for 10 weeks. n=12/group. **j-k.** Fecal non-esterified fatty acid (NEFA) (**j**) and total cholesterol (**k**) level from Aster-A^fl/fl^ (F/F) and littermates Aster-A^fl/fl^ CD4-Cre (A^ΔCD4^) mice fed *ad libitum* with HFD for 1.5 weeks. Statistical analysis: for panels a, repeated measure two-way ANOVA with Sidak’s multiple comparison test. For panel d, g, h, i, two-tailed unpaired Student’s t-test. For f, two-way ANOVA with Sidak’s multiple comparison test. ns, p>0.05, *p<0.05, **p<0.01, ***p<0.001, ****p<0.0001.

Loss of Aster-A in T cells also blunted the inflammatory adipose tissue changes characteristic of diet-induced obesity. Fat cell size was similar between genotypes but there was less macrophage infiltration in eWAT and iWAT of A^ΔCD4^ mice fed with HFD for 9 weeks (**Fig. 2e and S2d**). HFD-induced adipose expansion is associated with loss of embryonic adipose resident macrophages (ATM) and infiltration of a monocyte-derived inflammatory population (*31*, *32*). ATMs (F4/80^hi^CD11b^int^) were reduced in control mice 4 weeks post HFD, but this population was preserved in A^ΔCD4^ mice (**Fig. 2f**). A^ΔCD4^ mice also show decreased infiltration of inflammatory macrophages (F4/80^int^CD11b^hi^) at 9 weeks post HFD (**Fig. 2f**). Furthermore, after 9 weeks of HFD, A^ΔCD4^ mice showed reduced fat deposition in the liver (**Fig. 2g**) and reduced fasting glucose levels (**Fig. S2e**). They also maintained insulin and glucose sensitivity compared controls (**Fig. 2h-i**). These differences were not observed on normal chow diet (**Fig. S2f**). Thus, deletion of Aster-A from the T cell compartment alters systemic lipid metabolism in response to dietary fat intake.

Reduced adiposity in A^ΔCD4^ mice could not be attributed to differences in food intake (**Fig.S2g**). We also found similar energy expenditure and respiratory exchange ratios in both genotypes (**Fig.S2j-k**). To address the contribution of impaired dietary fatty acid uptake, we analyzed fecal composition from mice fed with HFD for 10 days. A^ΔCD4^ mice showed more non-esterified fatty acid (NEFA) in feces (**Fig. 2j**), despite similar fecal output, percentage of total nutrient absorbed, and content of unabsorbed fecal cholesterol (**Fig. 2k**, **S2j-k**). Collectively, our findings suggest that loss of Aster-A in T cells causes selective fatty acid malabsorption, leading to resistance to obesity and related metabolic pathologies.

### Loss of Aster-A in T cells enhances the Th17 signature in the small intestine

Given the evidence that loss of Aster-A in T cells impacted enterocyte responses to dietary fat, we sought to identify changes in SI resident T cell populations in A^ΔCD4^ mice. T-cell specific Aster-A deficiency did not impact thymocyte development or the homeostatic composition of peripheral T cells at steady state (**Fig. S3a-c**). We then performed transcriptional profiling of CD4^+^ and CD8^+^ T cells from jejunal lamina propria (LP) of mice that were acutely fed with olive oil (**Fig. S3d**). No meaningful transcriptional changes were observed between genotypes in the CD8^+^ T cells (**Fig. S3e**), however, a clade of genes was upregulated A^ΔCD4^ CD4^+^ T cells (**Fig. 3a**). Notably, Th17-associated transcripts were enriched, including those encoding the cytokines *Il17a, Il17f*, and *Il22*; the transcription factor *Rorc*, the Th17-specific chemokine receptors *Ccr6* and *Ccr7*, and other genes previously associated with gut Th17 function (**Fig. 3b**) (*33–36*). Gene ontology analysis of upregulated transcripts in A^ΔCD4^ T cells revealed enrichment in Th17 differentiation pathways (by KEGG), as well as positive effects on T cell differentiation (by GO) (**Fig. 3c**).

**Figure 3.**
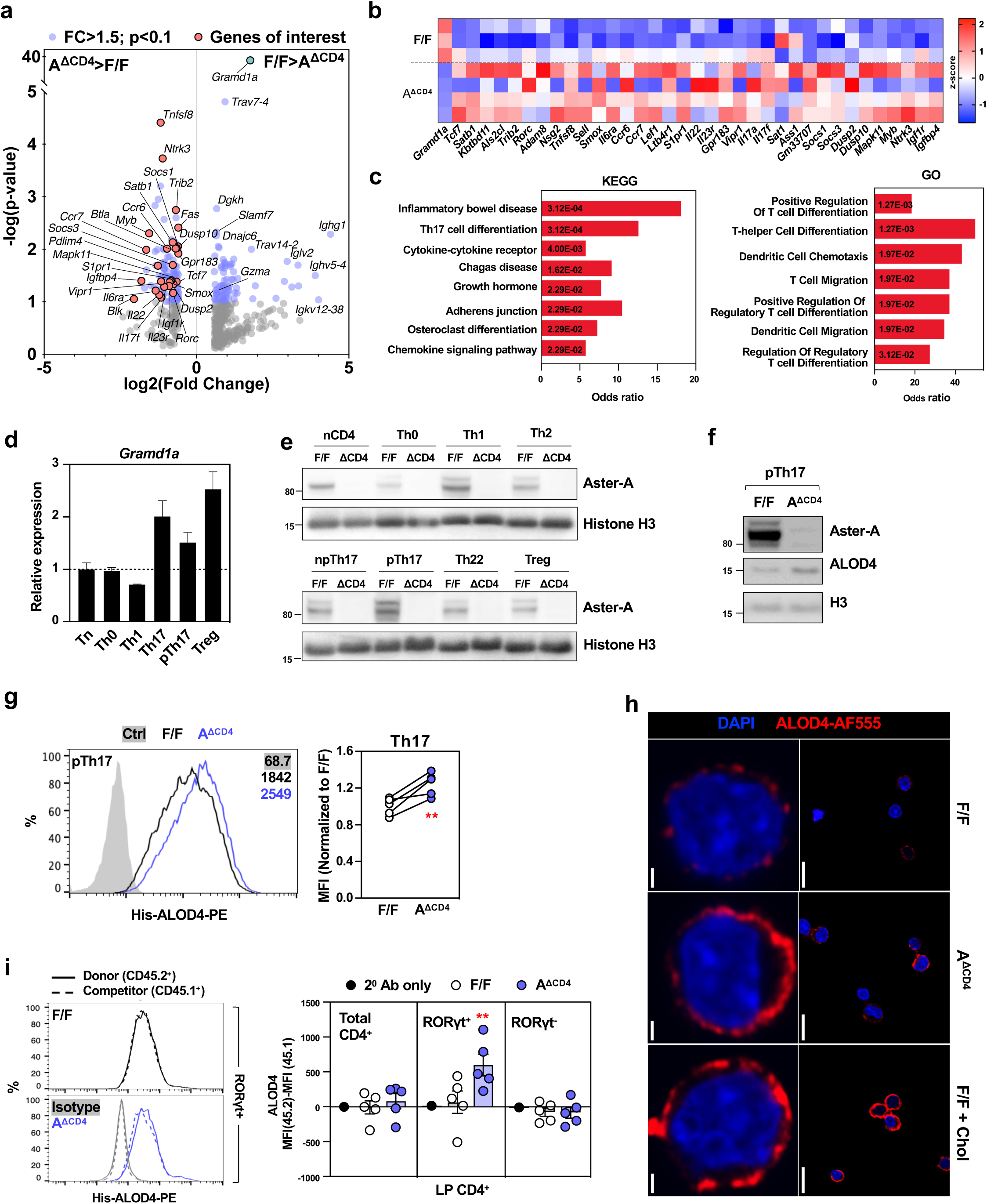
Aster-A maintains the homeostasis of PM accessible cholesterol in Th17 cells. **a.** Volcano plot showing differentially expressed genes in CD3^+^CD4^+^ lamina propria T cells from Aster-A^fl/fl^ (F/F) and littermates Aster-A^fl/fl^ CD4-Cre (A^ΔCD4^) mice gavaged with olive oil for 2 h. Blue dots indicate differentially expressed genes (FC>1.5, p<0.1). Red dots indicate genes of interest. n=3-4 biological replicates per group. **b.** Heat map of representative differentially expressed genes from (**a**). **c.** Gene set enrichment analysis of all differentially expressed genes in CD3^+^CD4^+^ lamina propria T cells comparing A^ΔCD4^ to F/F. **d-e.** mRNA (**d**) and protein (**e**) expression of Gramd1a (Aster-A) in *in vitro* differentiated Th cell subtypes. For western blot, same numbers of cells were used as input. npTh17: IL-6+TGFβ; pTh17: IL-6+TGFβ+IL-1β+IL-23. **f-g.** Western blot (**f**) and flow cytometry histogram (**g**) of ALOD4 in pTh17 cells (IL-6+TGFβ+IL-1β+IL-23) differentiated from Aster-A^fl/fl^ (WT) and Aster-A^fl/fl^ CD4-Cre (A^ΔCD4^) naive CD4^+^ T cells. (d) Mean fluorescence intensity (MFI) is shown. Secondary anti-His Tag-PE antibody alone, without His-ALOD4 staining was used as isotype control. **h.** ALOD4 staining (Red) of Aster-A^fl/fl^ (WT) and Aster-A^fl/fl^ CD4-Cre (A^ΔCD4^) pTh17 cells (IL-6+TGFβ+IL-1β+IL-23) or loaded with 25μM Methyl-β-cyclodextrin (MβCD)-Cholesterol for 1 h. Scale bar: 10μm (upper panels) and 1μm (lower panels). DAPI (Blue) staining indicate nucleus. **i.** (Left) histogram of accessible cholesterol levels (by ALOD4) in small intestine lamina propria RORγt^+^ T cells and (right) quantified difference of in total CD4^+^, RORγt^+^, or RORγt^-^ T cells, comparing Aster-A^fl/fl^ (WT CD45.2^+^) or Aster-A^fl/fl^ CD4-Cre (A^ΔCD4^ CD45.2^+^) to congenic competitor (CD45.1^+^) in mixed BM chimera. n=5 /group. Statistical analysis: for panel e, two-tailed Welsh’s t-test. For panel f, paired Student’s t-test. For panel i, two-way ANOVA with Tukey’s multiple comparison test. ns, p>0.05, **p<0.01, ****p<0.0001.

### Aster-A modulates the accessible cholesterol pool in Th17 cells

We next assessed how Aster-A modulates Th17 cell function. Among T helper cell subtypes in mice, Aster-A transcripts (*Gramd1a*) are most expressed in Th17 and Treg lineages in mice (**Fig. 3d**) and in Th17 cells in humans (**Fig. S4a**). Survey of published single cell RNA-sequencing data also demonstrated Gramd1a expression in human SI Th17 lineage (**Fig. S4b**). Aster-A protein was also abundant in the Th17 lineage (**Fig. 3e**). Aster-A mRNA and protein expression increased with Th17 differentiation (**Fig. S4d-e**). The preferential expression of Aster-A in Thj17 cells is consistent with the predicted binding of transcription factors that enforce Th17 lineage (including JUN, STAT3, EGR2, and SMAD4) at the promotor region of *Gramd1a* (**Fig. S4c**)(*37–39*).

We have shown previously that, in cells such as hepatocytes and enterocytes, Aster proteins mobilize the ‘accessible’ pool of cholesterol from plasma membrane (PM) to ER (*6*, *15*). Their function in lymphocytes, however, is undetermined. To test if Aster-A modulates PM cholesterol levels in T cells, we differentiated primary control or Aster-A-deficient (A^ΔCD4^) CD4 T cells to the Th17 lineage in a defined serum-free media (**Fig. S4e**) and used the ALOD4 probe to monitor accessible cholesterol levels on the PM (*3*). A^ΔCD4^ Th17 cells exhibited elevated accessible PM cholesterol compared to controls or to naïve T cells (**Fig. 3f-g**, **S4g**). Fluorescence microscopy suggested that accessible cholesterol was confined to ‘hot spots’ in control Th17 cell PM, but was more widely distributed in Aster-A deficient Th17 cells (**Fig. 3h**). This pattern was similar to that observed in T cells forcibly loaded with exogenous cholesterol (**Fig. 3h**). To examine intrinsic cholesterol homeostasis in the Th17 lineage *in vivo*, we generated mixed bone marrow chimeric mice using F/F control or A^ΔCD4^ bone marrow and congenic CD45.1^+^ competitor (**Fig. 3i**). We analyzed accessible cholesterol pool in lamina propria CD4 T cells by a combination of ALOD4 and immune marker staining (**Fig. S4h**). Comparison of the ALOD4 level of WT or A^ΔCD4^ to that of the competitor showed an increase in accessible PM cholesterol content in A^ΔCD4^ RORγt^+^ Th17 cells, but not in non-Th17 cells (**Fig. 3k**, **S4i**). Thus, Aster-A-mediated nonvesicular cholesterol transport maintains PM cholesterol homeostasis in Th17 cells.

### T cell receptor activation generates accessible PM cholesterol and recruits Aster-A

We next investigated the physiological signals that modulate accessible cholesterol abundance in T cells. We explored accessible cholesterol dynamics on Th17 cell PM after TCR activation by plate-bound anti-CD3/CD28. Control Th17 cells maintained constant accessible cholesterol levels over 4 hours, while Aster-A-deficient Th17 cells had elevated ALOD4 binding 1 and 2 hours post activation (**Fig. 4a**). Since Aster proteins are recruited to the PM by cholesterol in other cell types, we examined Aster-A localization during TCR activation. We expressed murine Aster-A tagged with mCherry at the N-terminus in EL4 cells. Aster-A was rapidly mobilized to the cell surface (marked by TCR staining) following TCR conjugation (**Fig. 4b**). We also used the OlyA probe to test whether Aster-A influenced the sphingomyelin (SM)-sequestered cholesterol pool (*4*), since SM and cholesterol concentrations correlate with TCR oligomerization and avidity (*23*). A^ΔCD4^ Th17 cells showed enhanced ALOD4 and Oly-A binding compared to WT Th17 cells at steady state, and both pools expanded upon TCR activation (**Fig. 5c**). This finding suggests that excess accessible PM cholesterol in Aster-A deficient cells funnels into the SM-sequestered pool.

**Figure 4.**
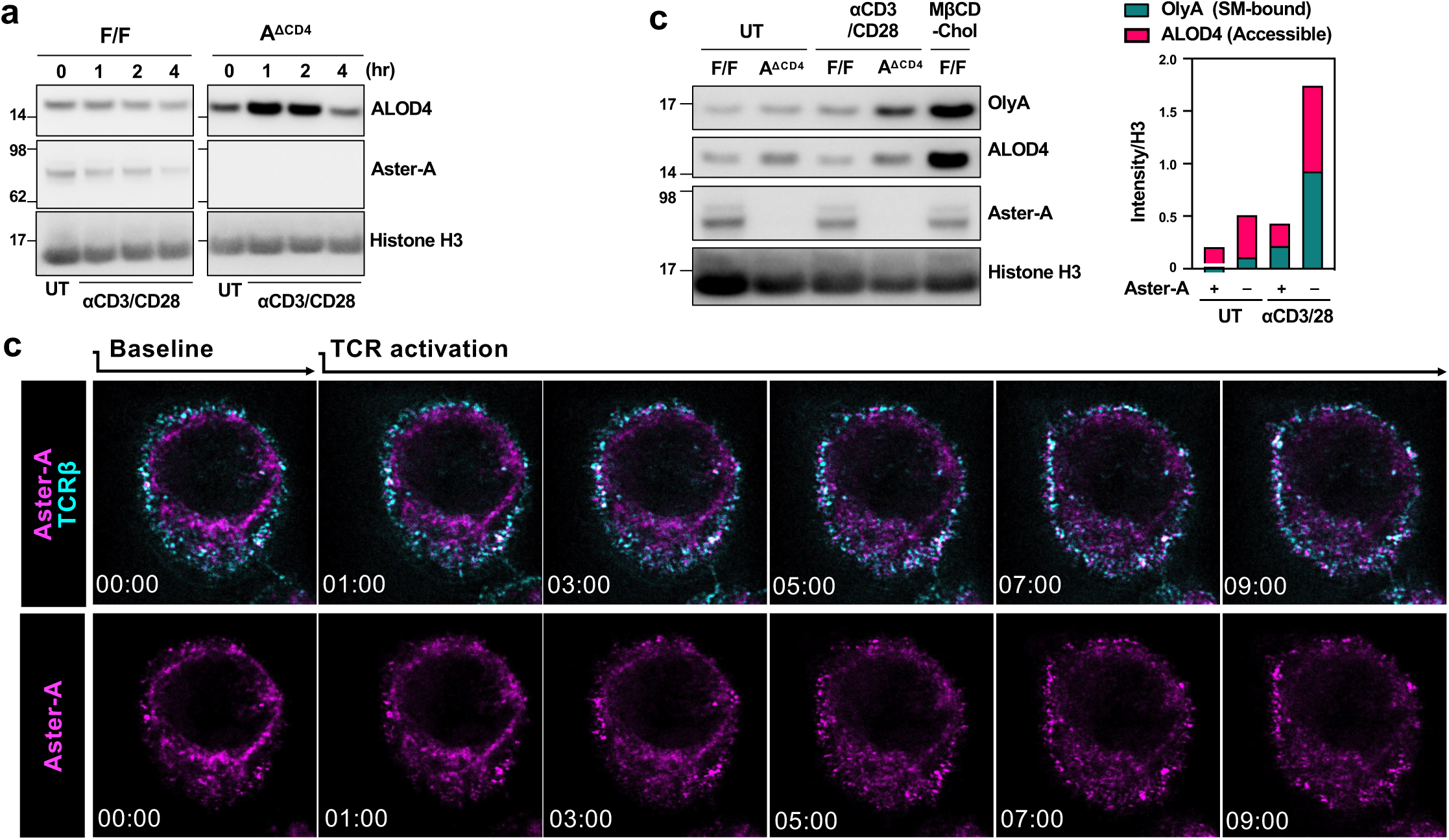
T cell receptor activation generates accessible cholesterol that rapidly recruits Aster-A to PM. **a.** Western blot analysis of ALOD4 staining (accessible cholesterol level) of in vitro differentiated Aster-A^fl/fl^ (WT) and Aster-A^fl/fl^ CD4-Cre (A^ΔCD4^) Th17 cells (IL-6+TGFβ+IL-1β+IL-23) that were reactivated with plate-bound 1μg/ml anti-CD3/CD28 antibody or PMA + Ionomycin for indicated times. **b.** Localization of Aster-A-mCherry in EL4 cells unstimulated, TCR crosslinked, or treated with 1μM Ionomycin. Plasma membrane TCRβ was stained with anti-TCRβ-AlexaFluor 647 antibodies. Scale bar: 2μm. **c.** Western blot analysis of ALOD4 (accessible cholesterol level) and OlyA (Sphingomyelin-sequestered cholesterol level) staining of in vitro differentiated Aster-A^fl/fl^ (WT) and Aster-A^fl/fl^ CD4-Cre (A^ΔCD4^) Th17 cells (IL-6+TGFβ+IL-1β+IL-23) that were reactivated with plate-bound 1μg/ml anti-CD3/CD28 antibody or loaded with 25μM MβCD-Cholesterol for 1 h.

**Figure 5.**
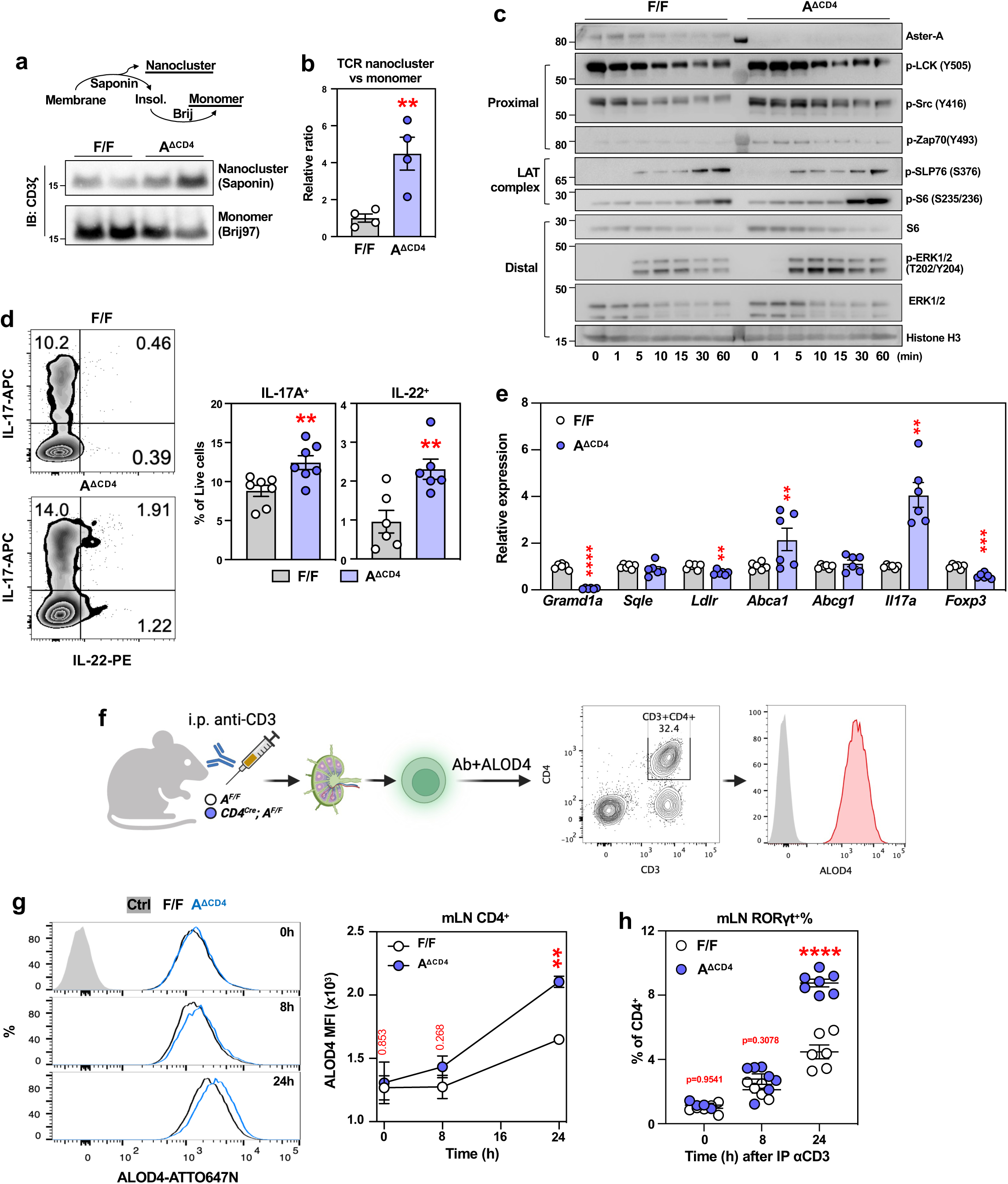
Aster-A regulates TCR nanoclustering and effector responses of Th17 cells through gating PM cholesterol level. **a-b.** Western blot analysis of TCR (CD3ζ) clustering in in vitro differentiated Aster-A^fl/fl^ (F/F) and Aster-A^fl/fl^ CD4-Cre (A^ΔCD4^) pTh17 cells. (**a**) Representative blot showing two biological replicates, and (**b**) quantification of Saponin-(nanocluster) versus Brij97-(monomer) extracted CD3ζ intensity, combined from two experiments, n=4 /group from 2 independent experiments. **c.** Western blot analysis of phosphorylation of LCK/Src, Zap70, SLP76, S6, and ERK1/2 following TCR activation. In vitro differentiated pTh17 cells were ‘rested’ overnight without differentiation cocktail or TCR stimulus and reactivated with plate-bound 1μg/ml anti-CD3/CD28 antibody for indicated times. Histone H3 as input control. Representative of 3 independent experiments. **d.** Frequency of *in vitro* differentiated pTh17 cells producing IL-17A or IL-22. Representative flow plot is shown on the left and quantifications on the right. n=6 /genotype, from two independent experiments. **e.** Gene expression from in vitro differentiated pTh17 cells as in (d). 4. **f.** Diagram of experimental set up to probe PM accessible cholesterol level during T cell activation *in vivo*. **g.** (left) Histogram of ALOD4 staining in mesenteric lymph node CD4+ T cells (CD45^+^CD3^+^CD4^+^) from Aster-A^fl/fl^ (F/F) and Aster-A^fl/fl^ CD4-Cre (A^ΔCD4^) mice that were intraperitoneally injected with anti-CD3ε (20μg/mouse) for indicated times or 20μg isotype control antibody (0h time point). (Right) Mean Fluorescence Intensity (MFI) of ALOD4 signal shown on the left. n=3-4 mice /genotype /time point, representative of two independent experiments. **h.** Frequency of RORγt^+^ population in mLN CD4^+^ T cells at indicated time points after anti-CD3ε injection. n=4-7 mice /group /time point, pooled from two independent experiments. Statistical analysis: for panel a, d, two-tailed Welsh’s t-test. For panel e, g, Welsh’s t-test with Holm-Sidak correction for multiple comparison. For h, two-way ANOVA with Sidak correction for multiple comparison. ns, p>0.05, *p<0.05, **p<0.01, ****p<0.0001.

### Aster-A restrains TCR nanoclustering and dampen proximal TCR signaling

PM cholesterol composition has been previously suggested to modulate membrane T cell receptor activity and avidity by impacting TCR nanoclustering (*40*). We asked if the accumulation of accessible and SM-sequestered PM cholesterol in A^ΔCD4^ Th17 cells (**Fig. 4a and 4c**) impacted TCR nanoclustering or signaling. Serial extraction of isolated Th17 membranes by saponin and Brij97 allowed separation of nanoclustered complexes from monomeric TCR (*41*). Quantification of CD3ζ chain abundance in these fractions indicated that the loss of Aster-A led to an increased TCR nanocluster to monomer ratio (**Fig. 5a-b**). Accordingly, following reactivation, A^ΔCD4^ Th17 cells exhibited enhanced proximal TCR signaling, as well as increased phosphorylation of S6 and ERK compared to controls (**Fig. 5c**, quantified in **Fig.S5a**). A^ΔCD4^ Th17 cells also showed increased frequency of effector cytokine (IL-17A and IL-22) production, had reduced Foxp3 expression (**Fig.5d-e**), and showed a moderate increase in proliferation (**Fig. S5b**). Of note, expression of genes linked to sterol synthesis and efflux were not altered in A^ΔCD4^ Th17 cells, suggesting that intracellular cholesterol homeostasis is largely maintained (**Fig. 5e**) (*42*). We conclude that Aster-A functions to restrain TCR nanoclustering and signaling in Th17 cells by removing PM cholesterol.

To test whether this phenomenon operates *in vivo*, we treated mice with intraperitoneal injection of CD3ε-specific antibody (**Fig. 5f**), an approach shown to elicit expansion and effector function of intestine-associated Th17 cells (*43*). Induction of the early activation marker CD69 in CD4^+^ T cells of mLN confirmed efficient TCR activation (**Fig. S5c**). Accessible cholesterol (by ALOD4) was unchanged in control mLN T cells at 8 h and moderately increased 24 h following anti-CD3ε injection (**Fig. 5g**). By contrast, A^ΔCD4^ mLN T cells accumulated accessible PM cholesterol at 8 h and this increased further by 24 h (**Fig. 5g**). Notably, anti-CD3ε treatment markedly expanded RORγt^+^ Th17 cells in the mLN of A^ΔCD4^ mice after 24 h, consistent with our finding that Aster-A dampens Th17 reactivation (**Fig. 5h**). Collectively these data demonstrate that TCR activation leads to increased PM accessible cholesterol, and that Aster-A is required for its timely removal to limit Th17 activation and expansion.

### Neutralization of IL-22 normalizes malabsorption in T-cell Aster-A deficiency

Lastly, we sought to understand the basis for malabsorption caused by gut Th17 cells in the absence of Aster-A. IL-22 is a mucosal Th17-associated cytokine in mice that maintains gut homeostasis and mucosal defense by increasing the production of anti-microbial peptides and promoting epithelium regeneration (*44–47*). Intriguingly, prior studies using stable overexpression or loss of IL-22 have also suggested a role in nutrient and energy metabolism (*48–53*). Consistent with our RNA-seq results (**Fig. 3a**), flow cytometry revealed increased IL-22^+^ frequency among A^ΔCD4^ LP T cells, and frequent co-expression with IL-17A (**Fig. 6a**). We further generated floxed-control and A^ΔCD4^ mice harboring an IL-22-GFP reporter allele (**Fig. S6a**) (*54*). We found more GFP^+^ cells in the LP CD4^+^ population from A^ΔCD4^ compared to control mice (**Fig. 6b**), while the GFP^+^ percentages of CD4^-^ T cells and innate lymphocytes were comparable (Fig. S6b). Induction of gut Th17 cells and IL-22^+^ populations in laboratory mice are known to depend on the colonization of Segmented Filamentous Bacteria (SFB) (*55*). In line with this, we found that the increased IL-22 response in A^ΔCD4^ mice was mainly attributed to TCR Vβ14^+^ T cells, a major clonal response to SFB antigen (**Fig. 6b**) (*56*, *57*). These findings confirmed that Aster-A deficient Th17 cells increased IL-22 producing frequency in the small intestine.

**Figure 6.**
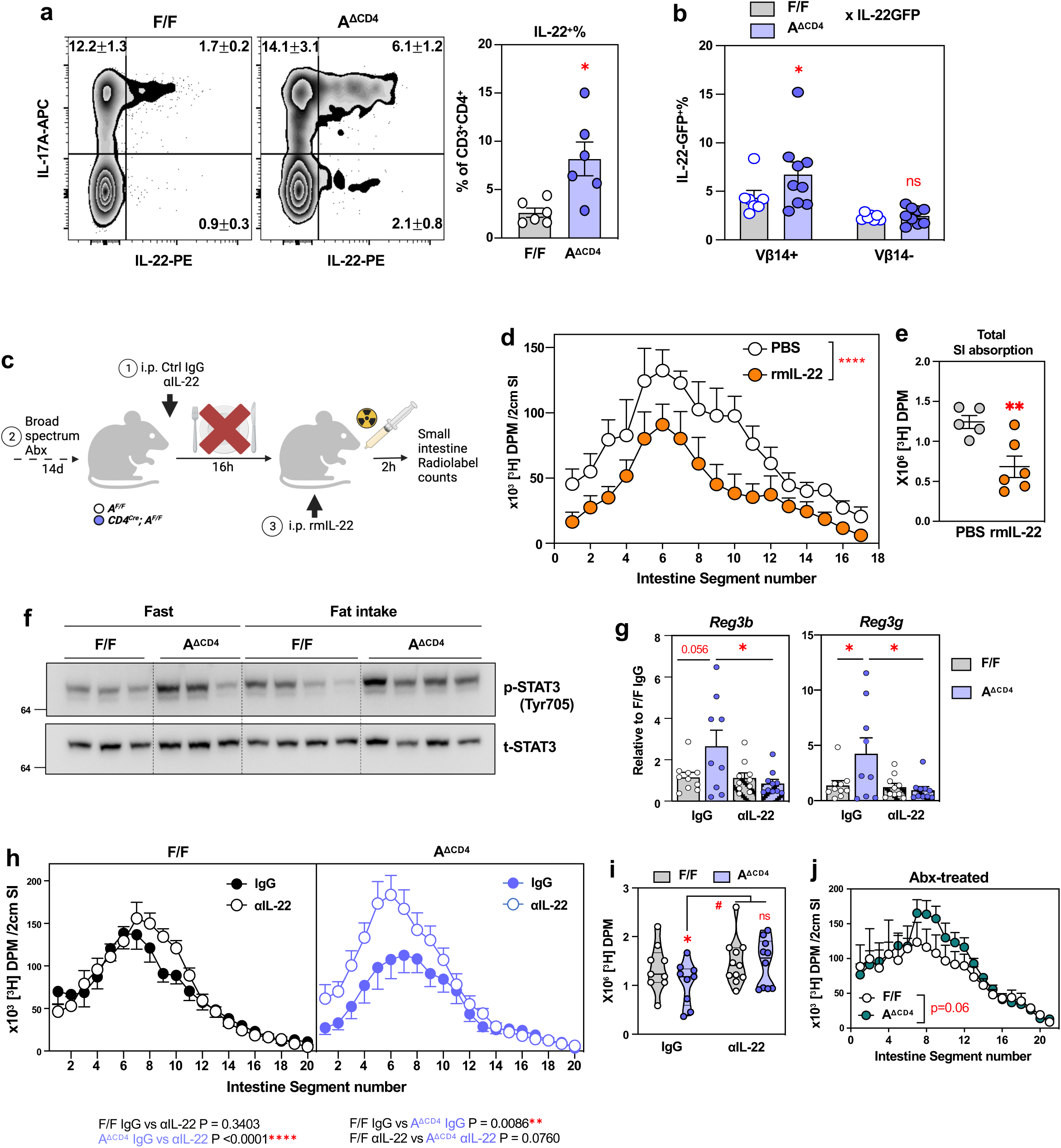
Aster-A deficiency leads to IL-22-mediated inhibition of fatty acid absorption. **a.** Percentage of IL-17A^+^ and IL-22+ T cells (CD45^+^CD90^+^CD3^+^CD4^+^) isolated from small intestine lamina propria of Aster-A^fl/fl^ (F/F) and littermates Aster-A^fl/fl^ CD4-Cre (A^ΔCD4^) mice gavaged with olive oil for 2 h. n=6 per group. **b.** Percentage of IL-22-GFP^+^ in Vb14^+^ or Vb14^-^ T cells (CD45^+^CD90^+^CD3^+^CD4^+^) from small intestine lamina propria of Aster-A^fl/fl^ Il22^GFP^(F/F x IL-22-GFP) and littermates Aster-A^fl/fl^ CD4-Cre Il22GFP (A^ΔCD4^ x IL-22-GFP) mice gavaged with olive oil for 2 h. n=7-9/group. **c.** Schematics for manipulation of IL-22 axis and absorption study. **d.** Distribution of radioactivity in intestinal segments of C57/BL6 wildtype mice after an oral challenge of olive oil containing [^3^H]-triolein (18:0) for 2 h. Mice were intraperitoneally injected with Vehicle (PBS) or 1.5μg IL-22 at 1 h prior to triolein challenge. n=5-6 /group. **e.** Total radioactivity in small intestines from experiments in **d**, (n=5/6 per group). **f.** Western blot analysis of phospho-STAT3 (Tyrosine 705) and total STAT3 (t-STAT3) in jejunal epithelial scrapings from Aster-A^fl/fl^ (F/F) and littermates Aster-A^fl/fl^ CD4-Cre (A^ΔCD4^) mice fasted for 16 h (Fast) or gavaged with olive oil (Fat intake) for 2 h. n=3-4 /group. **g.** *Reg3b* and *Reg3g* expression in terminal ileum of Aster-A^fl/fl^ (F/F) and littermates Aster-A^fl/fl^ CD4-Cre (A^ΔCD4^) mice intraperitoneally injected with anti-IL-22 (αIL-22) or control IgG antibody for 16 h as in (**c**). n=9-10 /group. **h-i.** Distribution of radioactivity in intestinal segments of Aster-A^fl/fl^ (F/F) and littermates Aster-A^fl/fl^ CD4-Cre (A^ΔCD4^) mice after an oral challenge of olive oil containing [^3^H]-triolein (18:0) for 2 h (**h**) and total absorption (**i**). Mice were treated with anti-IL-22 (αIL-22) or control IgG antibody for 16 h as indicated in (**c**). n=9-11 /group. **j.** Distribution of radioactivity in intestinal segments of Aster-A^fl/fl^ (WT) and littermates Aster-A^fl/fl^ CD4-Cre (A^ΔCD4^) mice after an oral challenge of olive oil containing [^3^H]-triolein (18:0) for 2 h. Mice were treated with broad-spectrum antibiotics in drinking water *ad libitum* (see methods) for 2 weeks. n=7-9 /group. Statistical analysis: for panel a, e, two-tailed Welsh’s t-test. For panel b, d, h, j, mixed-effects analysis. For g, two-way ANOVA with Tukey’s multiple comparison test. ns, p>0.05, #p<0.05, *p<0.05, **p<0.01, ****p<0.0001.

The receptor for IL-22 is a heterodimeric complex of IL-22R1 and IL-10R2 (*58*). Previous reports demonstrated that IL-22 acts on intestinal stem cells and secretory cells to facilitate epithelial regeneration and mucosal defense (*47*, *59–62*). We noted that publicly available single cell RNA-seq studies show co-expression of IL22R1/IL10R2 in mature absorptive enterocytes of the human small intestine (**Fig. S6c**) (*63*). Immunostaining further confirmed IL-22R1 localization to the apical membrane of jejunal villi (**Fig. S6d**). However, whether acute elevations in IL-22 levels are sufficient to suppresses fatty acid absorption is unknown. We treated mice with recombinant IL-22 for 1 hour prior to [^3^H]-triolein challenge (**Fig. 6c**). Exogenous IL-22 caused a marked reduction in label uptake across the length of SI (**Fig. 6d-e**, **S6e**), suggesting that IL-22 can acutely inhibit fatty acid uptake.

We next assessed signaling and transcriptional components downstream of IL-22R. IL-22R signaling, in particular through pSTAT3, induces transcription of antimicrobial proteins in SI (*46*, *64*). We found elevated phospho-STAT3 levels in isolated jejunal epithelial scrapings from A^ΔCD4^ mice (**Fig. 6f**), supporting the idea that IL-22 signals to jejunal epithelium, where majority of dietary fatty acid is absorbed (*65*, *66*). To directly test if elevated IL-22 contributed to fatty acid malabsorption in A^ΔCD4^ mice, we blocked IL-22 in by intraperitoneal administration of neutralizing antibodies prior to fasting (**Fig. 6c**). Control mice were treated with isotype IgG. A^ΔCD4^ mice showed elevated ileal expression of antimicrobial peptides *Reg3b* and *Reg3g*, consistent with enhanced IL-22R signaling, and this expression was blunted by blocking IL-22 (**Fig. 6g**). Moreover, neutralization of IL-22 ameliorated the impairment of fatty acid absorption in A^ΔCD4^ mice (**Fig. 6h-i**). Thus, loss of Aster-A in T cells inhibits fatty acid absorption largely through the effect of IL-22.

As the composition of the microbiome can be altered by IL-22 (*53*, *67*), we explored the contribution of microbiome to fatty acid malabsorption in A^ΔCD4^ mice. Control littermate and A^ΔCD4^ mice fed HFD had similar fecal bacterial composition by 16S sequencing (**Fig. S6f**). This observation is consistent with preserved IL-22 production from other immune compartments in A^ΔCD4^ mice (**Fig. S6b**). Furthermore, littermate mice that were individually housed or co-housed to deviate or normalize microbiota showed equal differences in HFD-induced weigh gain and total fat mass (**Fig. S6g-h**). Despite this, A^ΔCD4^ mice depleted of gut bacteria and fungus with a 2-week broad spectrum antibiotic regimen showed improvement from fatty acid uptake deficiency (**Fig. 6j**, **S6i-j**), and this effect correlated with loss of IL-22 from gut and mesenteric lymph nodes (**Fig. S6k**). We conclude that the effect of T-cell specific loss of Aster-A changes fatty acid absorption through direct T-cell-enterocyte communication mediated by IL-22.

## Discussion

The biophysical impact of PM cholesterol on T cell receptor assembly, clustering, and avidity has been well studied (*40*). These mounting evidence supports the concept that changes in PM cholesterol in T cells can have immunological consequences. However, the mechanisms that actively regulate T cell membrane cholesterol abundance, and thereby maintain signaling homeostasis, are poorly understood. Proximal T cell receptor dynamics are rapid, typically concluding within several minutes (*68*). Thus, it is reasonable to hypothesize that mechanisms must be in place to monitor and control PM cholesterol distribution to ensure TCR signaling quality. Here we have identified non-vesicular cholesterol transport as a metabolic checkpoint for Th17 activation and revealed that Aster-A is both a sensor and a responding rheostat for PM cholesterol dynamics during T cell activation. We further illustrated how membrane lipid homeostasis in immune cells can have broad impacts on systemic lipid metabolism, including on the control of dietary nutrient absorption.

Aster proteins are recruited to the ER-PM contact sites by excess membrane cholesterol and mobilize accessible PM for transport to the ER (*12*). We utilized the pool-specific probes ALOD4 (which detects accessible cholesterol), and OlyA (which detects sphingolipids-sequestered cholesterol) to visualize and quantify the dynamics of PM cholesterol distribution during T cell activation (*4*). We discovered that the accessible PM cholesterol pool expands immediately following TCR activation, leading to the recruitment of Aster-A. We also observed that some of this additional accessible cholesterol appeared to be funneled into the sphingomyelin-sequestered pool. Sphingomyelin and cholesterol-enriched domains, often interchangeably referred as lipid rafts, support TCR nanoclustering and reduce the threshold for activation (*22*, *23*). Accordingly, Aster-A-deficient resting Th17 cells exhibited an increased ratio of TCR nanoclusters to monomers, and this was associated with increased proximal TCR signaling. We posit that the heightened Th17 effector function observed *in vitro* and *in vivo* in the absence of Aster-A arises from both the inability to rapidly remove accessible cholesterol generated by TCR activation, and the increased TCR nanocluster predetermined by the chronic accumulation of sphingomyelin-cholesterol domains.

At steady state, the ER membrane contains ∼5% molar ratio of free cholesterol (*19*). Once excess cholesterol reaches the ER, it is rapidly converted to cholesterol esters by ACAT enzymes (*69*). Inhibition of ACAT1 activity has been shown to cause PM cholesterol accumulation and potentiate TCR activity in cytotoxic T cells (*26*, *70*), but the pathway that moved cholesterol from PM to ER to activate ACATs was heretofore unknown. Our observations in Th17 cells suggest that Aster-A most likely acts upstream of ACATs in this regulatory axis. Interestingly, loss of Aster-A does not seem to impact homeostatic T cell development and peripheral T cell composition. This is consistent with prior observations by us and others in which T cell cholesterol content was genetically perturbed by the loss of SCAP/SREBP2, ACAT1, or SULT2B1B (*22*, *26*, *71*). These studies suggest that effector T cells are particularly sensitive to changes in non-vesicular transport.

Combining the ALOD4 cholesterol probe with flow cytometry, we outlined the impact of non-vesicular cholesterol transport on Th17 homeostasis *in vivo*. We showed that accessible cholesterol accumulates in Aster-A-deficient lamina propria Th17 cells in a cell autonomous manner. This dynamic expansion of the accessible cholesterol pool also occurred following T cell activation by anti-CD3 *in vivo*, and it was associated with the expansion of Aster-A-deficient RORγt^+^ Th17 cells in mesenteric lymph nodes. These findings demonstrate that Aster-A is an indispensable guard of Th17 membrane homeostasis *in vivo*.

Hypercholesterolemia is linked to chronic inflammation. Excess LDL can be incorporated to membranes of immune cells and further promotes immunopathology (*72–76*). In particular, enhanced Th17-associated inflammation has been observed in various genetic- and diet-induced models of hypercholesterolemia (*77–82*). Recent studies have shown that vesicular- and non-vesicular cholesterol transport pathways converges at the PM (*83*, *84*). Cholesterol internalized by the LDLR receptor-mediated endocytosis is first liberated in the lysosome and then moves back to the PM where it can be mobilized to the ER by Asters (*19*, *84*). Our results strongly imply that Aster-A may be required to curtail hypercholesterolemia-induced inflammatory Th17 responses by resolving the PM accumulation of LDL-derived cholesterol. Further studies are required to fully understand potential anti-inflammatory roles of Asters in hypercholesterolemia and atherosclerosis.

Finally, our findings show that loss of membrane homeostasis in immune cells impacts physiological metabolism. Aster-A expression in T cells is required for appropriate fatty acid absorption and helps determine chronic metabolic responses to high fat diet. Aster-A deficiency led to increased Th17 activity in the small intestine, enhanced type-3 cytokine output, and impairs dietary fatty acid uptake. Neutralization experiments suggest that increased IL-22 production is the main mediator of this T-cell-intestinal crosstalk. Such a role for IL-22 in lipid and glucose metabolism is consistent with recent studies (*49–52*). Although Aster-A deficiency did not alter microbiome composition, broad spectrum antibiotics corrected the malabsorption of Α^ΔCD4^ animals. Interestingly, Aster-A deficiency led to increased IL-22 frequency in the SFB-associated clonal type, in line with the well-established role of SFB in inducing IL-22^+^ Th17 cells (*33*, *55*, *57*). These observations suggest that Aster-A tunes gut Th17 reactivity to commensal bacteria under homeostatic conditions. Our results, along with recent studies demonstrating the metabolic protective role of SFB-elicited Th17 cells (*52*), support the notion that mucosal lymphocytes exert important restraints on intestinal lipid handling.

## Acknowledgments

We thank all members of the Tontonoz, Tarling-Vallim, Mack, Young and Bensinger labs at UCLA for useful advice and discussions and for sharing reagents and resources. We also thank the UCLA CNSI Microscopy Core and UCLA Broad Stem Cell Flow Cytometry Core for their service and experimental suggestions. This study is supported by National Institutes of Health grant R01 DK126779 and Leducq Foundation Transatlantic Network of Excellence 19CDV04 to PT; Damon Runyon Foundation/Mark Foundation Postdoctoral Fellowship DRG-2424-21 and National Institutes of Health/National Center for Advancing Translational Sciences grant UL1TR001881 to YG; American Heart Association Postdoctoral Fellowship 903306 to JPK; American Heart Association Postdoctoral Fellowship 18POST34030388 to XX; National Institutes of Health grant CDI K12 07012023 to EW; American Diabetes Association Postdoctoral fellowship 1-19-PDF-043-RA to AF; National Institutes of Health grant T32 DK007180 and to AHN; National Institutes of Health grant F30 DK134050 to LFU.

---

**Figure S1.**
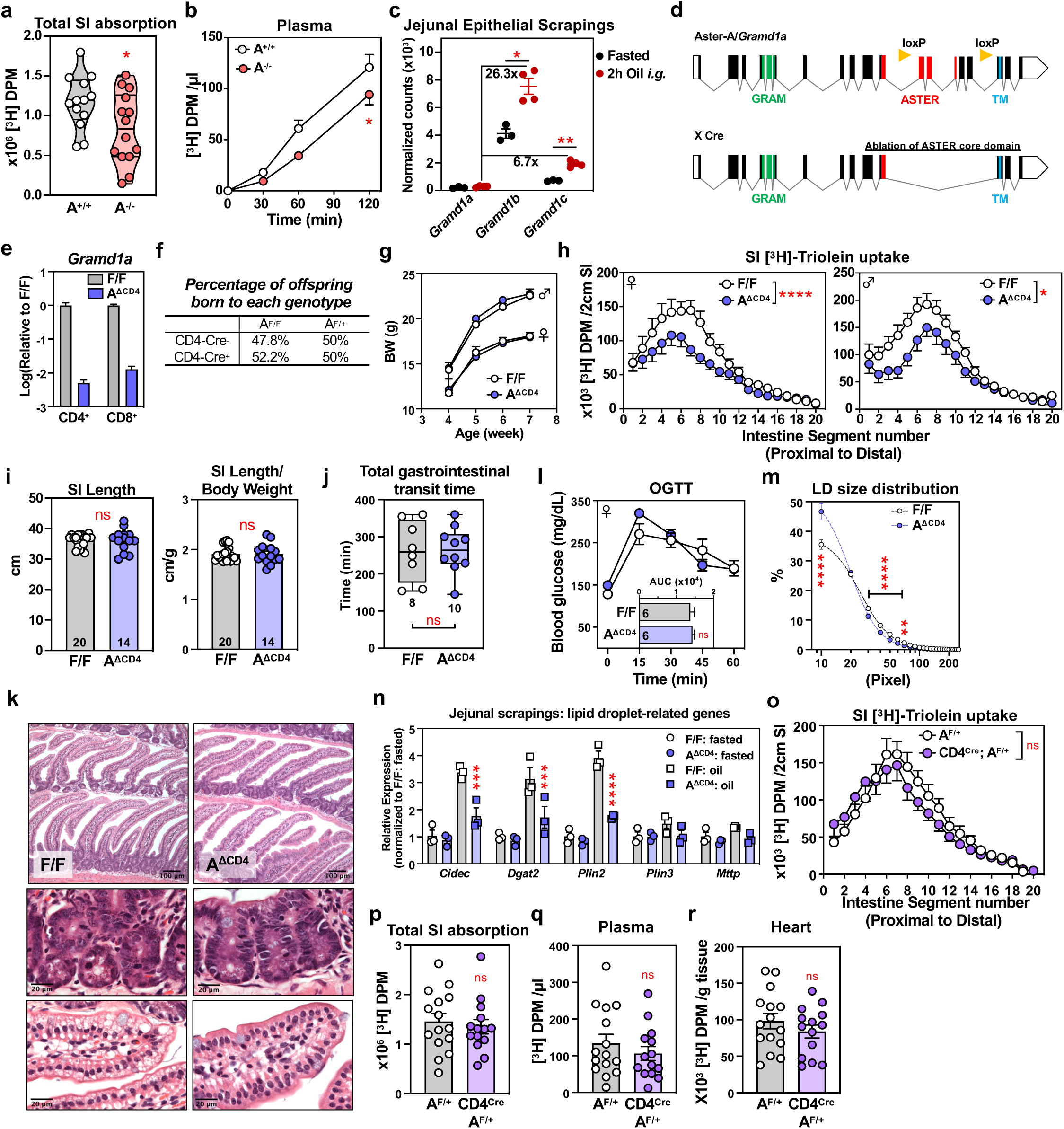
Loss of Aster-A in T cells leads to reduced intestinal fatty acid absorption without apparent pathology. **a.** Total radioactivity in small intestines from experiments in Figure 1a, n=13-14 /group. **b.** Kinetics of radioactivity in plasma of Aster-A wiltype (A^+/+^, n=13) and littermate global deficient (A^-/-^, n=14) mice after an oral challenge of olive oil containing [^3^H]-triolein (18:0). **c.** Expression of Aster-A, -B, -C (encoded by *Gramd1a, Gramd1b, Gramd1c*) in jejunal epithelium in wildtype mice fasted for 16 h or intragastrically gavaged (i.g.) with olive oil. **d.** Generation of Cre-driven Aster-A conditional KO mice **e.** mRNA expression of Aster-A (encoded by Gramd1a) in purified splenic CD4^+^ and CD8^+^ T cells. Expression values are normalized to Aster-A^fl/fl^ (F/F) and y-axis shown in log10 scale. **f.** Mendelian ratio of CD4-Cre^+^ and CD4-Cre^-^ progenies when crossed to Aster-A^fl/fl^ or Aster-A^fl/+^. **g.** Body weight of male and female Aster-A^fl/fl^ (F/F) and littermates Aster-A^fl/fl^ CD4-Cre (A^ΔCD4^) mice from 3 weeks (post weaning) to 7 weeks (early maturation). **h.** Distribution of radioactivity in intestinal segments of Aster-A^fl/fl^ (F/F) and littermates Aster-A^fl/fl^ CD4-Cre (A^ΔCD4^) mice after an oral challenge of olive oil containing [^3^H]-triolein (18:0) for 2 h. Figures are divided by female (left) and male (right), from same experiments in **Fig.1h**. **i.** Length (left) of small intestine of 8-12 week old Aster-A^fl/fl^ (F/F) and littermates Aster-A^fl/fl^ CD4-Cre (A^ΔCD4^) mice and length to body weight ratio (right). **j.** Total gastrointestinal transit time measured by carmine red assay. **k.** Representative hematoxylin and eosin (H&E) histology of jejunal section, villus tip, and crypt from Aster-A^fl/fl^ (F/F) and littermates Aster-A^fl/fl^ CD4-Cre (A^ΔCD4^) mice after an oral challenge of olive oil for 2 h. **l.** Small intestine glucose absorption measured by oral glucose tolerance test (OGTT). **m.** Distribution of lipid droplet (LD) size (in pixel) from experiments shown in **Fig.1n**. n=5 mice /group, combined from two independent experiments. **n.** mRNA expression of lipid droplet related gene in jejunal epithelial scrapings from Aster-A^fl/fl^ (F/F) and littermates Aster-A^fl/fl^ CD4-Cre (A^ΔCD4^) mice that were fasted for 16h or after an oral challenge of olive oil. **o-p.** Distribution of radioactivity in intestinal segments of male and female Aster-A^fl/+^ (n=15) and littermates Aster-A^fl/+^ CD4-Cre (n=14) mice after an oral challenge of olive oil containing [^3^H]-triolein (18:0) for 2 h (**o**) and total absorption (**p**). **q-r.** Radioactivity in plasma (**q**) and Heart (**r**) of Aster-A^fl/+^ (n=15) and littermates Aster-A^fl/+^ CD4-Cre (n=14) mice after an oral challenge of olive oil containing [^3^H]-triolein (18:0) for 2 h. Statistical analysis: for panels a, i, j, p, q, r, two-tailed Welch’s t-test. For b, c, l, m, repeated measure two-way ANOVA with Sidak’s multiple comparison test. For h, o, mixed-effects analysis. For n, two-way ANOVA with Tukey’s multiple comparison test. ns, p>0.05, *p<0.05, **p<0.01, ***p<0.001, ****p<0.0001.

**Figure S2.**
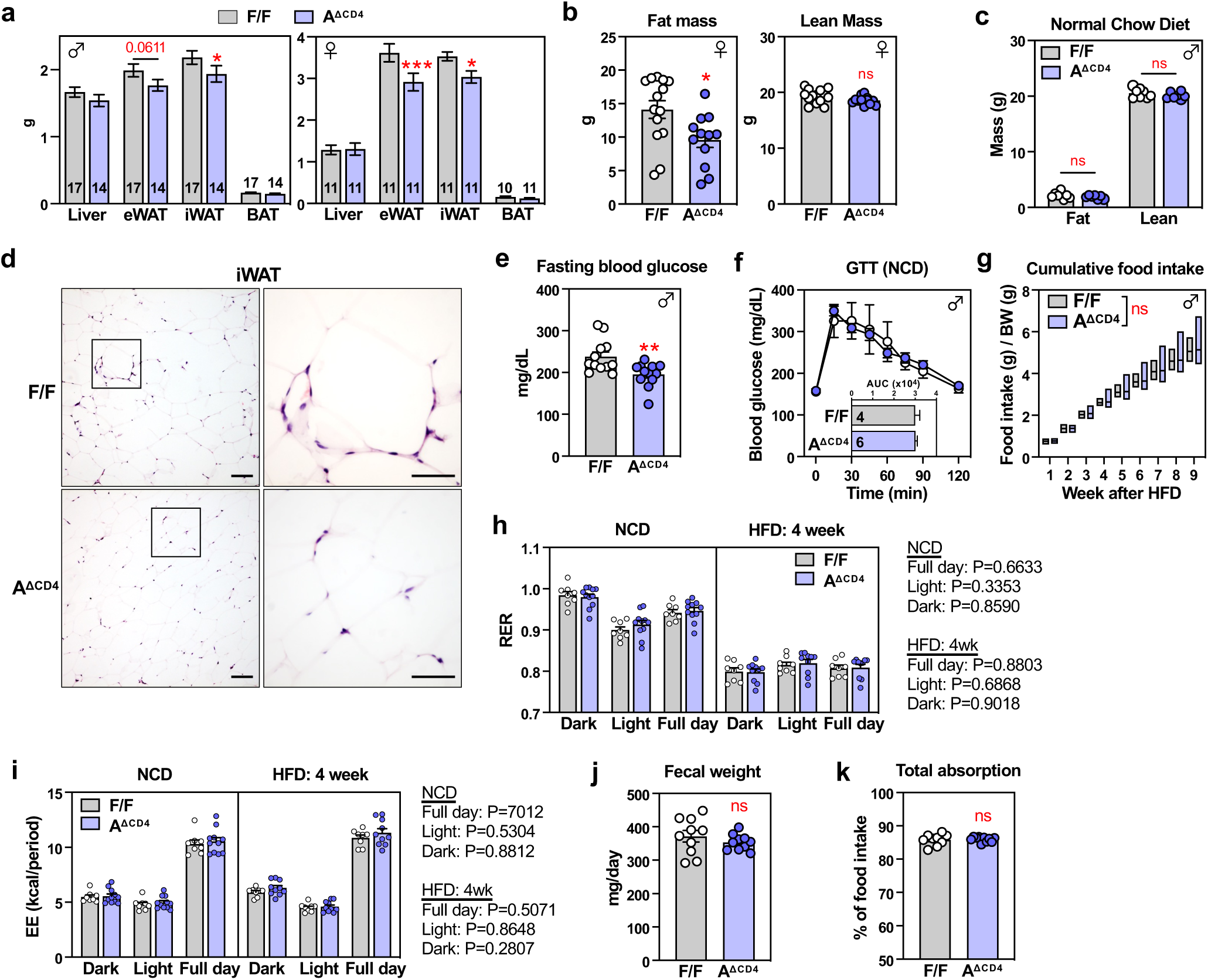
Impact of T cell-specific Aster-A deletion on diet-induced obesity. **a.** Weight of Liver, epididymal white adipose tissue (eWAT), inguinal white adipose tissue (iWAT), and brown adipose tissue (BAT) from male (left) and female (right) Aster-A^fl/fl^ (F/F) and littermates Aster-A^fl/fl^ CD4-Cre (AΔCD4) mice after 10 weeks of HFD feeding. **b.** Fat and lean mass of female mice after 9 weeks of HFD feeding determined by EchoMRI. n=14 (F/F) and 12 (A^ΔCD4^). **c.** Fat and lean mass of male mice in Fig.2 prior to HFD feeding. n=8 (F/F) and 7 (A^ΔCD4^). **d.** Representative hematoxylin and eosin (H&E) histology of inguinal WAT from mice in **Fig.2a**. Scale bar, 100μm; Inlay scale bar, 50μm. **e.** Fasting blood glucose level from male Aster-A^fl/fl^ (F/F) and littermates Aster-A^fl/fl^ CD4-Cre (A^ΔCD4^) mice fed *ad libitum* with 60 kcal% high fat diet (HFD) for 9 weeks. Mice were fasted for 4 h. n=12 /group. **f.** Glucose tolerance test (GTT) on 17-18 weeks old male Aster-A^fl/fl^ (F/F) and littermates Aster-A^fl/fl^ CD4-Cre (A^ΔCD4^) mice fed *ad libitum* with normal chow diet. n=4-6 /group. **g.** Cumulative food intake from single-housed male Aster-A^fl/fl^ (F/F) and littermates Aster-A^fl/fl^ CD4-Cre (A^ΔCD4^) mice fed *ad libitum* with 60 kcal% HFD. Weekly food intake was normalized to mouse body weight. n=10 /group. **h-i.** Respiratory exchange ratio (RER, **h**) and energy expenditure (EE, **i**) from male Aster-A^fl/fl^ (F/F) and littermates Aster-A^fl/fl^ CD4-Cre (A^ΔCD4^) mice fed *ad libitum* with normal chow diet or with 60 kcal% HFD for 4 weeks. fed or n=8-11 /group. **j.** Fecal output per day from single-housed male Aster-A^fl/fl^ (F/F) and littermates Aster-A^fl/fl^ CD4-Cre (A^ΔCD4^) mice fed *ad libitum* with 60 kcal% HFD for 1.5 weeks. n=10 /group. **k.** Total percentage of food absorption, as measured by (food intake-fecal weight)/food intake x 100, from mice in g. n=9-10 /group. Statistical analysis: for panels a, b, e, j, k, two-tailed Welch’s t-test. For panel b, mixed-effects analysis. For panel c, two-way ANOVA with Sidak’s multiple comparison test. For f-g, Repeated Measure two-way ANOVA with Sidak’s multiple comparison test. For h-i, ANCOVA with lean mass as a covariant. ns, p>0.05, *p<0.05, **p<0.01, ***p<0.001.

**Figure S3.**
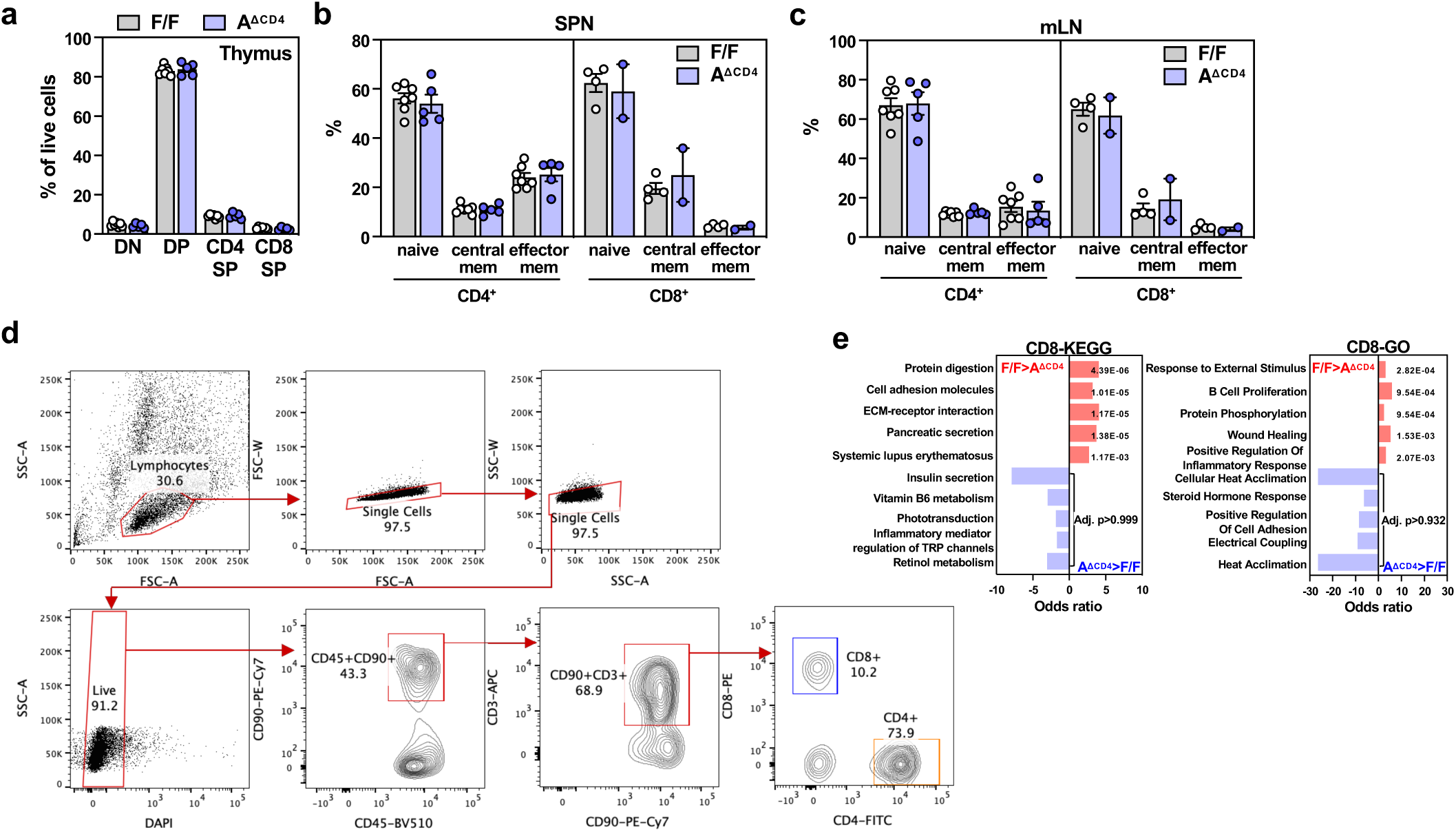
Aster-A deficiency does not impact peripheral T cell development or homeostasis in secondary lymphoid organs. **a.** Frequency of thymocytes in four major developmental stages. DN, CD8^-^CD4^-^; DP, CD8^+^CD4^+^; CD4 SP, CD4^+^CD8^-^; CD8 SP, CD8^+^CD4^-^. **b-c.** Frequency of naive (CD62L^+^CD44^-^), central memory (CD62L^+^CD44^+^), effector memory (CD62L^-^CD44^+^) T cells in spleen (**b**) or mesenteric lymph nodes (mLN, **c**) CD4+ and CD8+ T cells. **d.** Gating strategy for FACS sorting of lamina propria CD4^+^ and CD8^+^ T cells. **e.** Gene set enrichment analysis of all differentially expressed genes in CD3^+^CD8α^+^ lamina propria T cells comparing A^ΔCD4^ to F/F.

**Figure S4.**
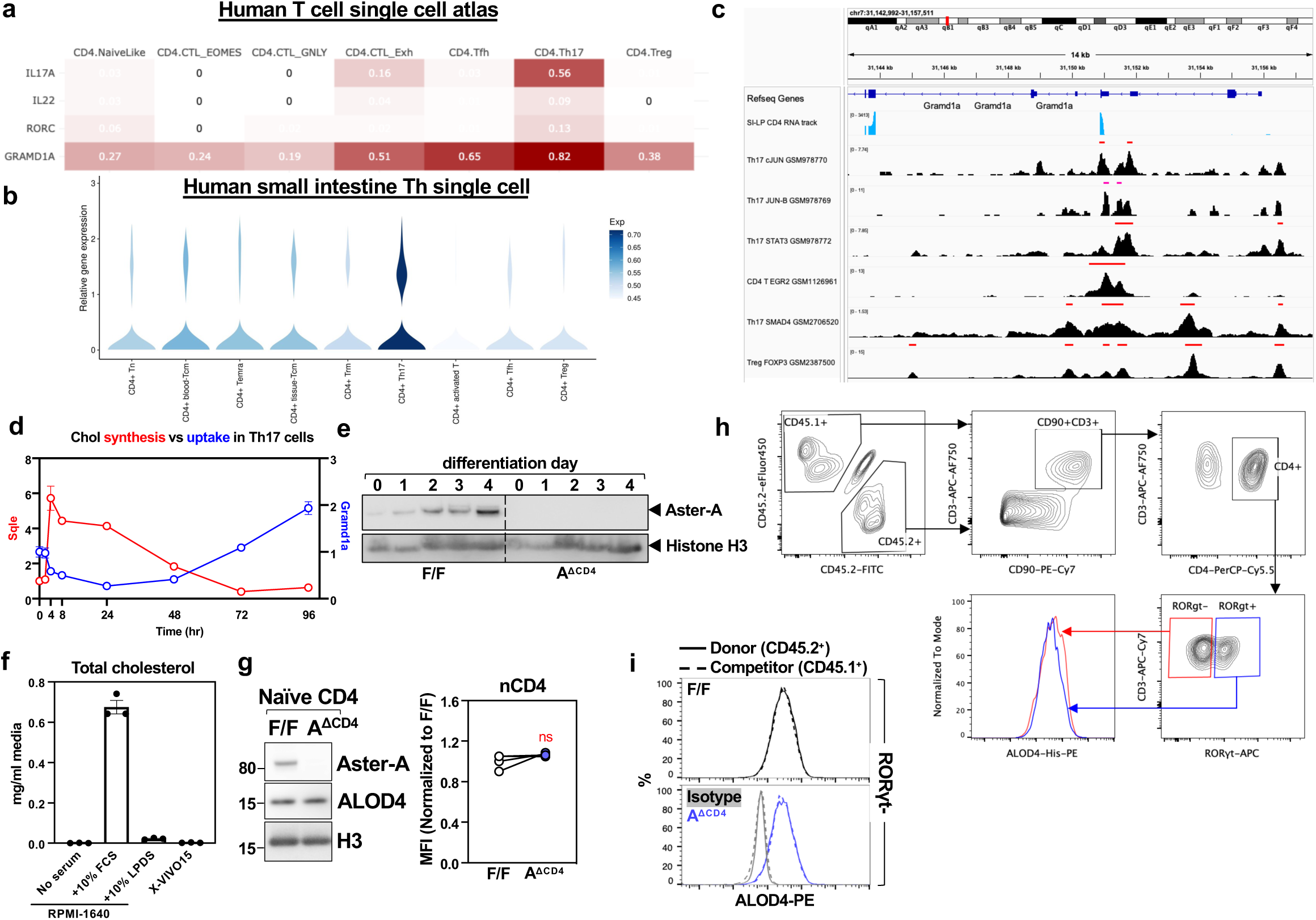
Aster-A maintains membrane cholesterol homeostasis in Th17 cells. **a-b.** Expression of IL17A, IL-22, RORC, GRAMD1A from human T cell single cell atlas (Swiss Portal for Immune Cell Analysis, **a**) and small intestine T helper (Th) cell single-cell analysis (scIBD database, **b**). **c.** mRNA sequencing track of *Gramd1a* proximal region from small intestine lamina propria (SI-LP) T cells, aligned with chromatin immunoprecipitation sequencing track of cJUN, JUN-B, STAT3, EGR2, SMAD4, and FOXP3 in Th17 cells from Cistrome database. **d.** Expression of SREBP2 target *Sqle* compared to *Gramd1a* in different time points of naive T cell to Th17 cell differentiation, determined by RT-qPCR. **e.** Protein level of Aster-A in time points of naive T cell to Th17 cell differentiation. **f.** Concentration of cholesterol in standard RPMI media (Std media), standard media supplemented with 10% fetal bovine serum or liproprotein deficient serum, or X-VIVO15. **g.** ALOD4 level by western blot (left) or flow cytometry (right) comparing naive Aster-A^fl/fl^ (F/F) and Aster-A^fl/fl^ CD4-Cre (A^ΔCD4^) CD4 T cells. **h.** Gating strategy for distinguishing congenic RORγt^+^ T cells from experiment in Fig. 3i. **i.** Histogram of accessible cholesterol levels (by ALOD4) in small intestine lamina propria RORγt^-^ T cells comparing Aster-A^fl/fl^ (F/F CD45.2^+^) or Aster-A^fl/fl^ CD4-Cre (A^ΔCD4^ CD45.2^+^) to wildtype congenic competitor (CD45.1^+^). Statistical analysis: for panel f, paired Student’s t-test. ns, p>0.05.

**Figure S5.**
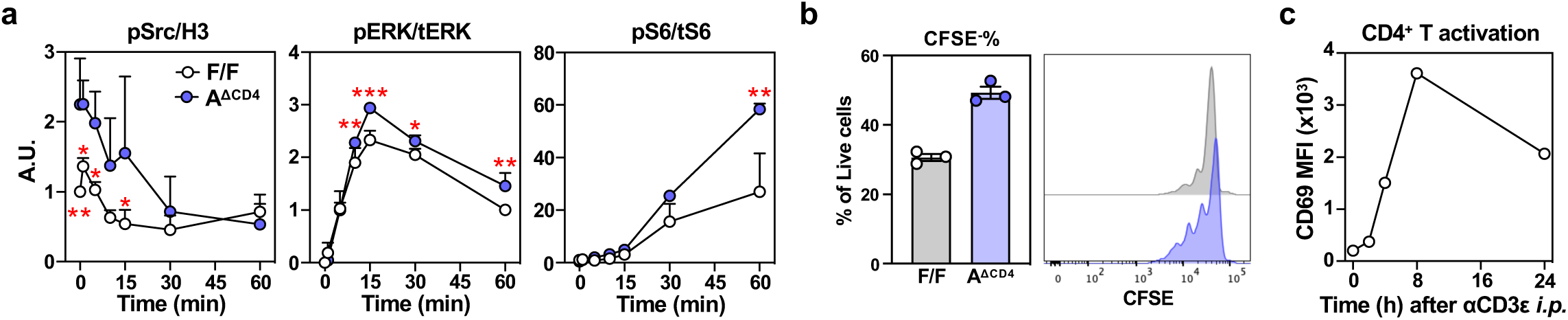
Aster-A restrains TCR signaling and proliferation of Th17 cells. **a.** Quantified band intensity from TCR signaling blots as in **Fig.5c**, from 2-3 independent experiments. **b.** Proliferation index comparing F/F or A^ΔCD4^ T cells and co-cultured competitor (CD45.1^+^) T cells under Th17 differentiation condition. n=3. **c.** Frequency of CD69^+^ in mesenteric lymph node CD4^+^ T cells in C57/BL6 mice injected with 20μg anti-CD3ε antibody for indicated times. Statistical analysis: for panel a, repeated measures two-way ANOVA. *p<0.05, **p<0.01, ***p<0.001.

**Figure S6.**
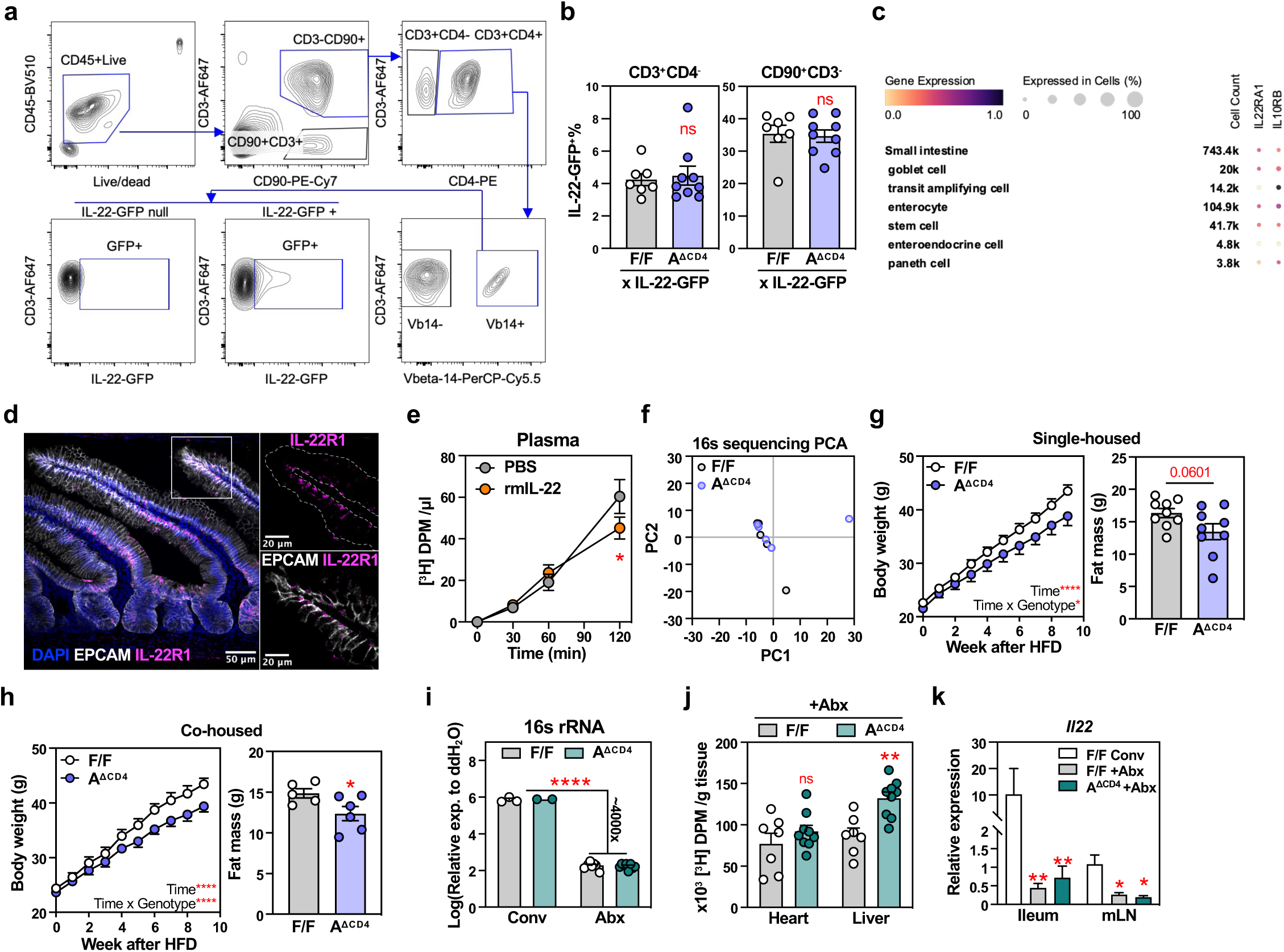
Impact of lymphocyte-specific Aster-A on dietary fat uptake is dependent on IL-22 but independent of microbiota alterations. **a.** Gating strategy for identifying IL-22-GFP^+^ populations from lamina propria lymphocytes, related to Fig. 6b. **b.** Frequency of IL-22-GFP^+^ in CD3^+^CD4^-^ (non-CD4 T cells) or CD90^+^CD3^-^ (ILCs), from small intestine lamina propria of Aster-A^F/F^ *Il22*^GFP^(WT x IL-22-GFP) and littermates Aster-A^fl/fl^ CD4-Cre *Il22*^GFP^ (A^ΔCD4^ x IL-22-GFP) mice gavaged with olive oil for 2 h. **c.** IF microscopy of IL-22R1 (magenta) in jejunum. EPCAM (white) marks basolateral surface and DAPI (blue) marks nucleus. **d.** Gene expression of IL-22 receptor subunits (IL22RA1 and IL10RB) in major small intestinal epithelial cell types. Expression data were obtained from prior human single cell RNA sequencing experiments deposited in the Cell x Gene database. **e.** Kinetics of radioactivity in plasma of C57/BL6 wildtype mice after an oral challenge of olive oil containing [^3^H]-triolein (18:0) for 2 h. Mice were intraperitoneally injected with PBS or 1.5ug recombinant mouse IL-22 (rmIL-22) at 1 h prior to triolein challenge. n=5-6 /group. **f.** Principal component analysis (PCA) of microbial Operational Taxonomic Unit abundance from 16S rRNA sequencing. Littermte Aster-A^fl/fl^ (F/F) and Aster-A^fl/fl^ CD4-Cre (A^ΔCD4^) male mice were single-housed from 8 weeks old for 2 weeks before feces collection. n=5 /group. **g.** Body weight (left) of littermates Aster-A^fl/fl^ (F/F) and Aster-A^fl/fl^ CD4-Cre (A^ΔCD4^) male mice that were single housed from 5-week-old and fed with HFD from 8-week-old for 9 weeks. Fat mass measured by EchoMRI is shown on the right panel. n=9 /group. **h.** Body weight (left) of littermates Aster-A^fl/fl^ (F/F) and Aster-A^fl/fl^ CD4-Cre (A^ΔCD4^) male mice that were mix co-housed and fed with HFD from 8-week-old for 9 weeks. Fat mass measured by EchoMRI is shown on the right panel. n=5-6 /group. **i.** 16s rRNA qPCR in fecal DNA from mice treated with broad-spectrum antibiotics (Abx) in drinking water ad libitum for 2 weeks as in **Fig.6j**. n=2-9 /group. **j.** Radioactivity in heart and liver from antibiotics treated Aster-A^fl/fl^ (F/F) and littermates Aster-A^fl/fl^ CD4-Cre (A^ΔCD4^) after an oral challenge of olive oil containing [^3^H]-triolein for 2 n=7-9 /group. **k.** mRNA expression of *Il22* in terminal ileum or mesenteric lymph nodes from mice in (**i-j**). n=4-9 /group. Statistical analysis: for panel b, two-tailed Student’s t-test. For panel e, repeated-measures two-way ANOVA with Sidak’s multiple comparison test. For panel g and h, mixed-effects analysis (for weight gain curve) and two-tailed Student’s t-test (for fat mass). For panel i-j, two-way ANOVA with Tukey’s multiple comparison test. For k, Fisher’s LSD test. *p<0.05, **p<0.01, ****p<0.001.

